# Two independent DNA repair pathways cause mutagenesis in template switching deficient *Saccharomyces cerevisiae*

**DOI:** 10.1101/2023.06.25.546467

**Authors:** Yangyang Kate Jiang, Eleanor Medley, Grant W. Brown

## Abstract

Upon DNA replication stress, cells utilize the post-replication repair pathway to repair single-stranded DNA and maintain genome integrity. Post-replication repair is divided into two branches: error-prone translesion synthesis, signaled by PCNA mono-ubiquitination, and error-free template switching, signaled by PCNA poly-ubiquitination. In *Saccharomyces cerevisiae*, Rad5 is involved in both branches of repair during DNA replication stress. When the PCNA poly-ubiquitination function of Rad5 is disrupted, Rad5 recruits translesion synthesis polymerases to stalled replication forks, resulting in mutagenic repair. Details of how mutagenic repair is carried out, as well as the relationship between Rad5-mediated mutagenic repair and the canonical PCNA-mediated mutagenic repair, remain to be understood. We find that Rad5-mediated mutagenic repair requires the translesion synthesis polymerase ζ but does not require other yeast translesion polymerase activities. Furthermore, we show that Rad5-mediated mutagenic repair is independent of PCNA binding by Rev1 and so is separable from canonical mutagenic repair. In the absence of error-free template switching, both modes of mutagenic repair contribute additively to replication stress response in a replication timing-independent manner. Cellular contexts where error-free template switching is compromised are not simply laboratory phenomena, as we find that a natural variant in *RAD5* is defective in PCNA poly-ubiquitination and therefore defective in error-free repair, resulting in Rad5- and PCNA-mediated mutagenic repair. Our results highlight the importance of Rad5 in regulating spontaneous mutagenesis and genetic diversity in *S. cerevisiae* through different modes of post-replication repair.

---

Genetic information must be accurately passed to the next generation to ensure cell survival. DNA replication stress is a barrier to high-fidelity genome transmission (Zeman and Cimprich 2014; Saxena and Zou 2022), and if not resolved in a timely manner can result in accumulation of single-stranded DNA (ssDNA), mutagenesis, and other forms of genome instability. One of the pathways that cells utilize to maintain genome integrity is the post-replication repair (PRR) pathway, which bypasses damaged bases and repairs ssDNA gaps formed at replication forks. PRR is mediated in part by post-translational modifications of the proliferating cell nuclear antigen (PCNA) (Gao et al. 2017; Arbel et al. 2020). In *Saccharomyces cerevisiae*, PCNA K164 is mono-ubiquitinated by the Rad6-Rad18 complex in response to ssDNA revealed by DNA replication stress or encounters between the replisome and damaged bases. Mono-ubiquitinated PCNA (PCNA-Ub) recruits error-prone translesion synthesis (TLS) DNA polymerases to mediate error-prone repair. PCNA-Ub can be poly-ubiquitinated by the Rad5-Ubc13-Mms2 complex, which signals for error-free template switching events where the nascent sister chromatid is used as a repair template (Hoege et al. 2002; Minca and Kowalski 2010). Independent of PCNA poly-ubiquitination, Rad5 can also recruit error-prone polymerases directly to stressed replication forks (Xu et al. 2016; Gallo et al. 2019). Therefore, both error-prone and error-free pathways can repair ssDNA gaps and bypass base lesions to promote cell survival, although with entirely different mutational outcomes.

*S. cerevisiae* has three TLS polymerases: Rev1, Pol η (Rad30), and the Pol ζ complex (Rev3, Rev7, Pol31, and Pol32). An inserter-extender model has been proposed for the TLS process where one TLS polymerase inserts an incorrect base at the site of damage, and another TLS polymerase extends beyond the lesion until normal replication by the more accurate replicative polymerases resumes. In this model, Rev1 acts as a scaffold protein to recruit Rad30 as the inserter, followed by Pol ζ which acts as the extender (Rizzo and Korzhnev 2019). Intriguingly, depending on the context of the DNA damage, Rev1 may also serve as an inserter (Zhou et al. 2010; Wiltrout and Walker 2011; Wang and Xiao 2020). While the error-prone TLS polymerases are implicated in bypassing DNA lesions, several lines of evidence indicate roles for TLS polymerases in repair events that do not involve DNA damage. DNA replisome mutants incur TLS polymerase-dependent mutations (Northam et al. 2006; Aksenova et al. 2010; Becker et al. 2014; Denkiewicz-Kruk et al. 2020), as do strains depleted of dNTPs (Northam et al. 2010; Gallo et al. 2019). Thus, replication stress in the form of DNA base lesions, and lesion-less replication stress induced genetically or chemically, are both overcome by the action of error-prone DNA polymerases, promoting cell survival at the cost of mutagenesis.

TLS polymerases are called into play during DNA replication stress, and PCNA-Ub and Rad5 both provide platforms for recruiting TLS polymerases to stressed DNA replication forks. We previously inferred the existence of two distinct mutagenic PRR pathways, because a *rad5* mutant lacking template switching activity has a higher mutation rate than cells that lack *rad5* entirely (Gallo et al. 2019), but direct support for two mutagenic pathways is lacking. Here we test the hypothesis that Rad5 and PCNA-Ub mediate distinct mutagenic repair pathways during DNA replication stress. We identify the TLS polymerase catalytic and scaffolding activities that are required for mutagenic repair and probe the relationship between mutagenic repair and replication timing. Finally, we show that a natural *RAD5* variant that is linked to increased mutagenesis is defective in error-free repair, providing a biological context where increased Rad5 mutagenic repair promotes genetic diversity.

## Materials and Methods

### Yeast strains and media

Yeast strains (Reagent Table) used in the study were derived from BY4741 (Brachmann et al. 1998) with the exception of Y8800 and Y8930 which were derived from W303. Strains were constructed using standard yeast genetic and molecular cloning methods and were cultured under standard conditions. The *rev3-D1142A-D1144A* (*rev3-cd*) and *rad5-E783D-I791S* (*rad5-RM*) strains were constructed by homology-directed repair with annealed synthetic oligos as repair templates co-transformed with the CRISPR-Cas9 plasmid pUB1306 (constructed by Gavin Schlissel in Jasper Rine’s lab, and a kind gift from Dr. Elçin Ünal) expressing sgRNA sequences targeting the respective genes. The *6HIS-POL30* strain was constructed using the same approach, but with a PCR product from YIplac128-His6-POL30 (Davies and Ulrich 2012) as the repair template. Expression of all *RAD5, REV3*, and *REV1* alleles was confirmed by immunoblot analysis (Figure S1).

### Hydroxyurea sensitivity assay

Saturated cultures were diluted to OD_600_ = 0.5 and diluted serially from 1:10 to 1:1,000 in a 96-well plate. 4 µL of each dilution was spotted onto YPD plates containing the indicated concentrations of hydroxyurea (HU). Plates were incubated at 30°C for three days prior to imaging.

### Growth rate measurement

Saturated cultures were diluted to OD_600_ = 0.05 in 100 µL YPD in a 96-well plate. Plates were incubated in a Tecan Sunrise microplate reader at 30°C with agitation, and OD_600_ measurements were taken every 15 minutes for 48 hours. Generation time was calculated using the growth rate algorithm designed by Danielle Carpenter (https://scholar.princeton.edu/sites/default/files/bot-steinlab/files/growth-rate-using-r.pdf). Three technical replicates were performed for each of two independent experiments (n = 6). Statistical support for growth rate differences was assessed with a 2-sided t-test.

### Mutation rate assay

Mutation rates at *CAN1* or *URA3* were measured using a Luria-Delbrück fluctuation test (Lang and Murray 2008; Lang 2018). For each fluctuation test, saturated cultures were diluted 1:10,000 in 10 mL of fully supplemented SD media, and 96 30 µL cultures were grown in a flat-bottom 96-well plate. After overnight incubation, 4 to 6 cultures were pooled to calculate N(t) using a hemocytometer. For the remaining 90 to 92 cultures, 220 µL of SD-arginine+60 mg/L canavanine was added to each well. The plate was then sealed with an adhesive plastic plate seal and incubated for 3 days. Wells were scored for colony formation, and mutation rate was calculated according to the Poisson distribution (Lang 2018). Statistical support for mutation rate differences was assessed with a 2-sided t-test.

For the mutation rates with the *URA3* reporter (Figure 4), saturated cultures were diluted 1:2,500 in 10 mL SD-all media, and 96 30 µL cultures were grown. N(t) was determined as indicated above. Cultures were spotted on SD + 50mg/L uracil + 0.1% 5-fluoroorotic acid plates, incubated for 3 days prior to scoring for growth, and the mutation rate was calculated according to the Poisson distribution (Lang 2018). Statistical support for mutation rate decreases was assessed with a 1-sided t-test.

**Figure 4.**
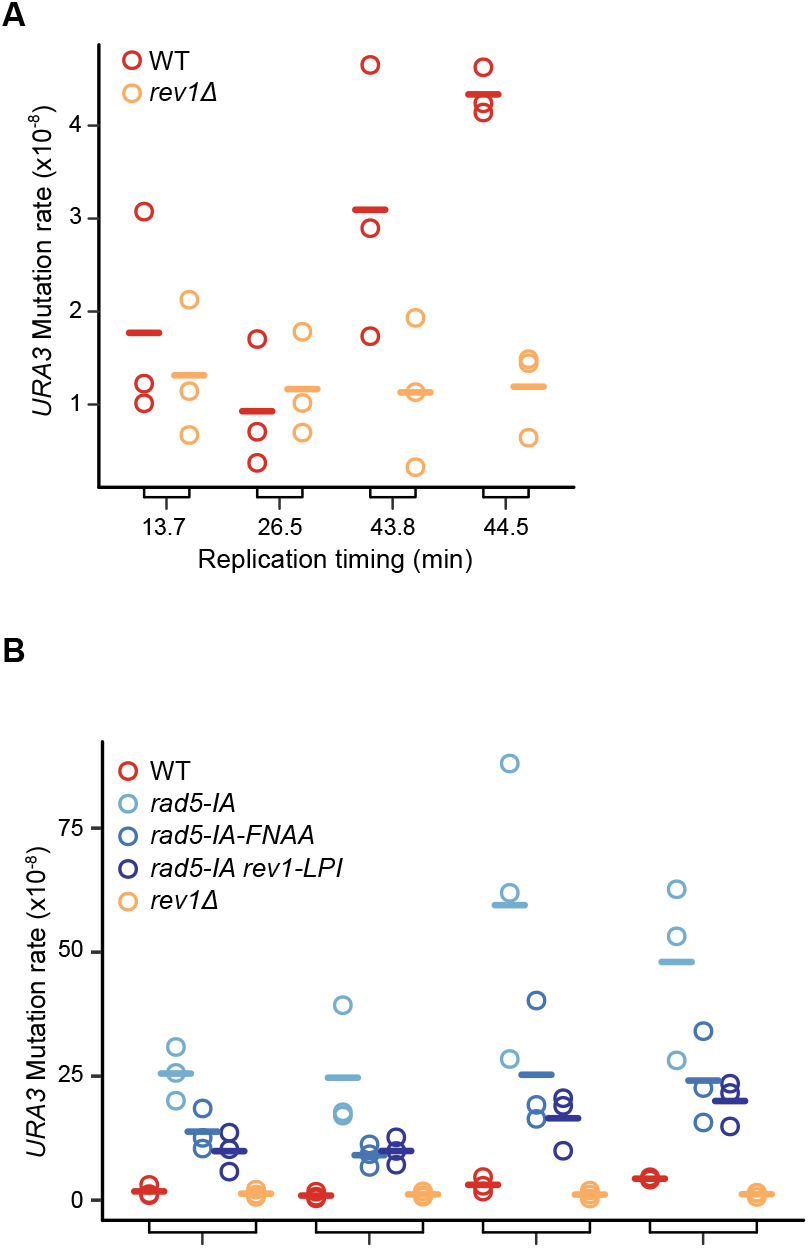
Rad5- and PCNA-mediated error-prone repair act independent of replication timing. (A-B) Spontaneous mutation rates at *URA3* inserted at four different positions on chromosome VI are plotted for the indicated strains. The replication timing is indicated for each *URA3* insertion. Each replicate (n = 3) is plotted, and solid bars represent the means.

### Fluorescence microscopy

RPA foci formation was assessed as previously described (Gallo et al. 2019), using Rfa1-GFP. Briefly, strains were grown to mid logarithmic phase in YPD, fixed with paraformaldehyde, washed and resuspended in mounting medium containing 4′,6-diamidi-no-2-phenylindole (DAPI). Images were acquired by illuminating samples with a Lumencor Spectra X Light Engine with 395/25 excitation filter (for DAPI) and 470/24 excitation filter (for GFP) and taking eleven 0.4 µm z-stack images on a Nikon Eclipse Ti2 inverted microscope using the brightfield, GFP (513/44 emission filter) and DAPI (430/35 emission filter) channels. Image stacks were converted to a single image by projecting the maximum pixel value at each pixel position in CellProfiler version 3.1.9 (McQuin et al. 2018), and foci formation and budding index were scored by visual inspection in ImageJ (https://imagej.nih.gov/ij/). At least 100 cells were scored for each replicate. Statistical support for differences in RPA foci positive cells was assessed with a 2-sided t-test.

### Flow cytometry

Rnr3-GFP strains were grown to mid logarithmic phase in YPD, and 1mL of the culture was pelleted and resuspended in 1mL PBS. The cells were briefly sonicated and analyzed on BD FACS Canto II or BD LSR Fortessa with the 488nm laser and 530/30 filter. A total of 10,000 events were collected and gated for Rnr3 positive events in FlowJo (BD Life Sciences). Statistical support for differences in Rnr3-GFP positive cells was assessed with a 2-sided t-test.

### Ubiquitination analysis

PCNA tagged with 6x His was affinity-purified under denaturing conditions as described (Davies and Ulrich 2012). In brief, ∼70 OD units of cells in mid logarithmic phase were collected and treated with 240 mM sodium hydroxide/1% 2-mercaptoethanol followed by 6.35% trichloroacetic acid. Precipitated proteins were resuspended in 6M guanidine hydrochloride and incubated with 25 µL of Ni-NTA agarose beads (Qiagen) overnight. The beads were washed and proteins were eluted in loading buffer containing 8 M urea, 200 mM Tris-HCl (pH 6.8), 1 mM EDTA, 5 % SDS, 0.1 % bromophenol blue, 1.5 % dithiothreitol. The eluate was heated at 65°C for 10 minutes prior to fractionating on 10% Tris-glycine SDS-PAGE. PCNA and ubiquitinated PCNA were detected by immunoblotting using a polyclonal rabbit anti-PCNA antibody (a generous gift of Dr. Marco Muzi-Falconi, used at 1:2,000 dilution) and a monoclonal mouse anti-ubiquitin antibody (Cell Signaling Ubiquitin (P4D1) Mouse mAb, #3936, 1:1,000 dilution).

### Yeast whole cell extract preparation and analysis

For validation of protein expression (Figure S1), strains were grown to mid logarithmic phase and 5-10 OD units of cells were collected and diluted with YPD to a total volume of 9 mL. One mL of 100% trichloroacetic acid was added to the culture followed by 15 minutes at room temperature on a rocking platform. The culture was then centrifuged at 2,000 rpm for 5 minutes and the pellet was washed with 1 mL of 1M HEPES buffer (pH 7.5). The pellet was resuspended in 50 µL 1x SDS-PAGE loading buffer (50 mM Tris-Cl (pH 6.8), 100 mM dithiothreitol, 2% SDS, 0.1% bromophenol blue, 10% glycerol) and lysed with 0.5 mm glass beads by vortexing at 4°C for 3-10 minutes. An additional 50 µL 1x SDS-PAGE loading buffer was added to the sample and the extract was incubated at 95°C for 5 minutes prior to gel loading. Proteins tagged with 6His10FLAG were detected by immunoblot-ting using an anti-FLAG mouse antibody (Sigma ANTI-FLAG M2 antibody F3165, 1:5,000 dilution), and Pgk1 was detected with an anti-Pgk1 mouse antibody (Invitrogen PGK1 monoclonal antibody, #459250, 1:10,000 dilution).

### Yeast 2-hybrid assay

Yeast 2-hybrid (Y2H) assay was performed as previously described (Sing et al. 2018). *RAD5* and *UBC13* were amplified from genomic DNA with primers containing attB sites and inserted into pDONR201 or pDONR221 (Invitrogen) vectors by Gateway BP reactions according to the manufacturer’s instructions. The resulting entry vectors containing the ORFs were then cloned into Y2H destination vectors that encode the Gal4 activation domain (pDEST-AD) or the Gal4 DNA-binding domain (pDEST-DB) (Vidal 2000) by Gateway LR reactions according to the manufac-turer’s instructions, resulting in fusion of the Gal4 domains to the N-termini of the proteins. The activation domain or DNA-binding domain plasmids were then transformed into Y8930 (*MATα*) or Y8800 (*MAT***a**) strains, respectively. To test for protein-protein interactions, strains were mated on YPD plates, diploids were selected on SD-Trp-Leu plates, and grown to saturation in SD-Trp-Leu media. Saturated cultures were diluted to OD = 2 in a 96-well plate in 200 µL water and serial diluted from 1:4 to 1:64. 4 µL of each dilution was spotted on SD-Trp-Leu or SD-Trp-Leu-His, and plates were incubated for three days at 30°C prior to imaging.

### Statistical analysis

Statistical analysis and data visualizations were performed in R 4.1.1 (https://www.r-prsoject.org/). F-tests were performed to identify samples with unequal variance, and Student’s t-tests or Welch’s unequal variance t-tests were performed accordingly.

### Data availability

Strains and plasmids are available upon request. The authors affirm that all data necessary for confirming the conclusions of the article are present within the article, figures, tables, and supplemental materials.

## Results

### Rad5-mediated error-prone repair requires Pol ζ

We previously established the presence of Rad5-mediated TLS polymerase activity on undamaged ssDNA templates during DNA replication stress using an I916A mutant of Rad5 (*rad5-IA*) (Figure 1A) (Gallo et al. 2019). I916A is within the RING finger domain of Rad5 and disrupts the interaction between Rad5 and Ubc13. The *rad5-IA* mutant is therefore incapable of PCNA poly-ubiquitination (Ulrich 2003; Toth et al. 2022) and so lacks error-free (template switching) repair activity. In agreement with our previous study, we observed that *rad5-IA* has wild type HU sensitivity (Figure 1B), but has elevated spontaneous and HU-induced mutation rates (Figure 1C; p = 0.02 for spontaneous, p = 0.01 for HU-induced), indicating that *rad5-IA* relies on error-prone repair to resist replication stress caused by HU (Gallo et al. 2019). Hence, we used *rad5-IA* as a genetic model to study the mechanisms of error-prone repair during DNA replication stress.

**Figure 1.**
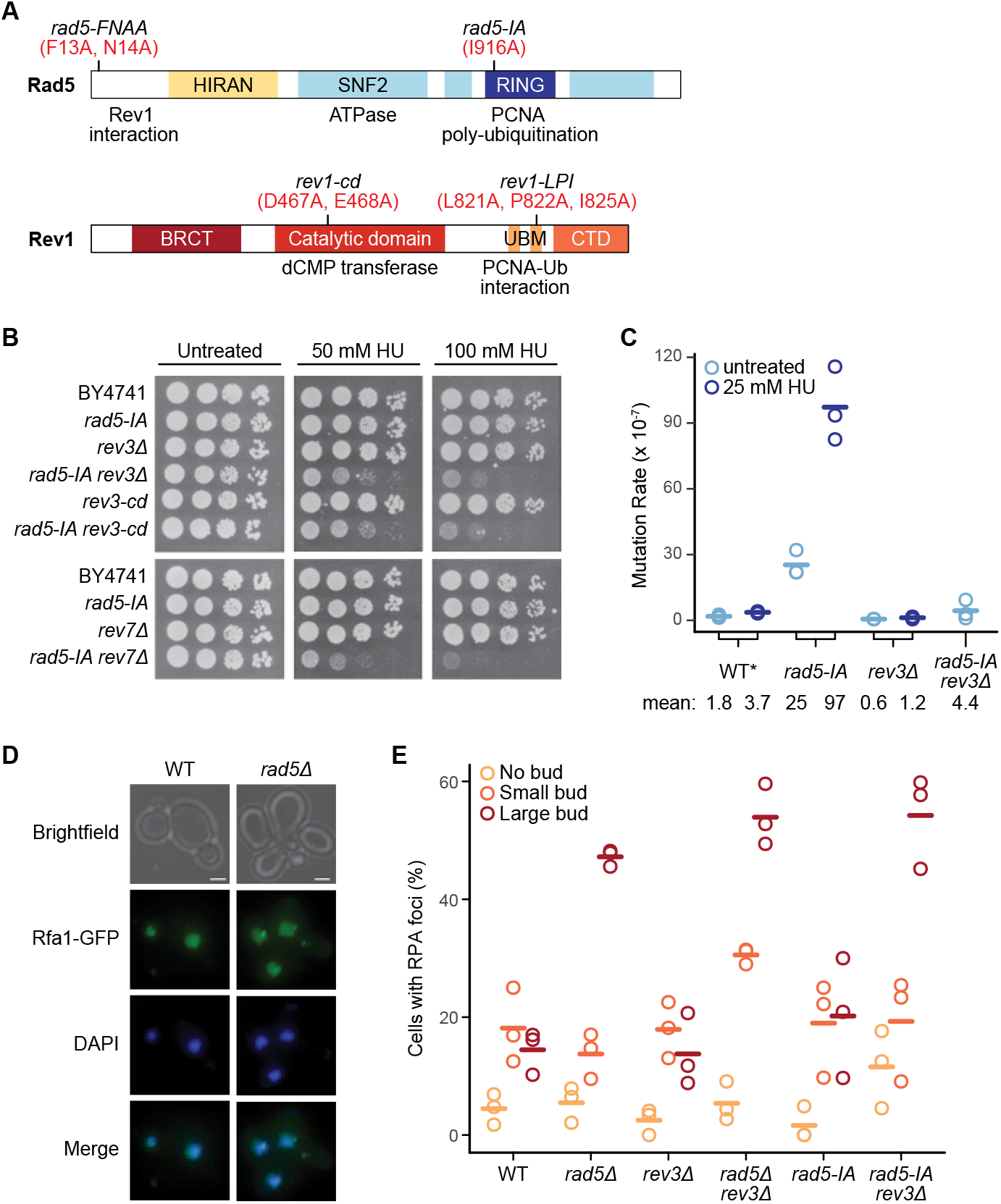
Rad5-mediated error-prone repair requires the catalytic activity of Pol ζ. (A) Schematics of the domain structures of Rad5 and Rev1. Conserved domains are indicated, as are the positions of the amino acid substitution mutants used in this study (in red). The names of the alleles corresponding to each mutant are indicated above each schematic, and the function of each domain is indicated below. Note that the SNF2/ATPase domain in Rad5 is tripartite. (B) HU sensitivity assay of the indicated strains. Cultures were serially diluted and spotted on solid media containing the indicated concentrations of HU. Plates were incubated for three days prior to imaging. (C) Spontaneous and HU-induced mutation rates of the indicated strains are plotted. WT* (DGY19) carries *RAD5* tagged with 6His10FLAG, as *rad5-IA* is 6His10FLAG-tagged. Each replicate (n = 3) is plotted, and solid bars represent the means, which are also indicated below the plot. (D) Fluorescence microscopy of RPA focus formation. Representative images of wild type and *rad5Δ* strains are shown. Cells are visible in the brightfield images, diffuse nuclear RPA and RPA foci are shown (Rfa1-GFP), and nuclei were stained with DAPI. Scale bar, 2 μm. (E) Quantification of RPA focus formation in the indicated strains. At least 100 cells were counted for each strain, grouped based on the budding status, and the percentage of cells with RPA foci for each group is plotted. Each replicate (n = 3) is plotted, and solid bars represent the means.

Error-prone repair in *rad5-IA* requires the TLS gene *REV1* (Gallo et al. 2019). Since Rev1 can recruit other TLS polymerases through its C-terminus to carry out DNA repair (Rizzo and Korzhnev 2019), we asked whether additional TLS polymerases (Rad30 and DNA polymerase ζ (Pol ζ)) are involved in Rad5-mediated error-prone repair. Deletion of the Pol ζ catalytic subunit *REV3* in *rad5-IA* led to increased HU sensitivity (Figure 1B) and decreased mutation rate (Figure 1C; p = 0.01), indicating that Pol ζ is required for error-prone repair in *rad5-IA*. Consistent with Pol ζ being required, deletion of the Pol ζ regulatory subunit *REV7* or the disruption of Rev3 catalytic activity using a catalytically dead mutant (Siebler et al. 2014) both resulted in increased HU sensitivity of *rad5-IA* (Figure 1B). Absence of error-prone repair in *rad5*Δ and *rad5-IA rev1Δ* strains results in accumulation of ssDNA that can be detected by measuring nuclear foci of the ssDNA binding protein RPA (Figure 1D and (Gallo et al. 2019)). We observed an increase in the percentage of cells with RPA foci in the large budded (G2/M) cells of *rad5-IA rev3Δ* relative to *rad5-IA* (Figure 1E; p = 0.01), indicating that Rev3 is required to prevent accumulation of ssDNA in *rad5-IA*. Taken together, our data support the conclusion that the catalytic activity of Pol ζ is required for the error-prone repair of ssDNA tracts that accumulate in *rad5-IA* during endogenous and HU-induced replication stress.

### Error-prone repair of ssDNA does not require Rad30 or the catalytic activity of Rev1

We next asked whether Rad30 is involved in error-prone repair in *rad5-IA*. Deletion of *RAD30* in *rad5-IA* did not change the HU sensitivity or the mutation rate, suggesting that Rad30 is not responsible for mutagenic repair in *rad5-IA* (Figure 2A and 2B). Since the catalytic activity of Rev1 plays a role in defective-replisome-induced mutagenesis in the absence of damage (Northam et al. 2014), we asked if the dCMP transferase domain of Rev1 plays a role in error-prone repair in *rad5-IA* using a mutant allele encoding catalytically inactive Rev1 (*rev1-cd*) (Figure 1A, (Zhou et al. 2010)). The HU sensitivity of *rad5-IA rev1-cd* was similar to wild-type both in plate-based assays and in quantitative growth rate measurements, in contrast to the high HU sensitivity of *rad5-IA rev1Δ* (Figure 2A and 2C). Moreover, disrupting the catalytic activity of Rev1 in *rad5-IA* did not change either the spontaneous mutation rate or the HU-induced mutation rate (Figure 2B). Collectively, these data suggest that mutagenic repair in *rad5-IA* does not involve the dCMP transferase activity of Rev1, either during endogenous replication stress or during HU treatment.

**Figure 2.**
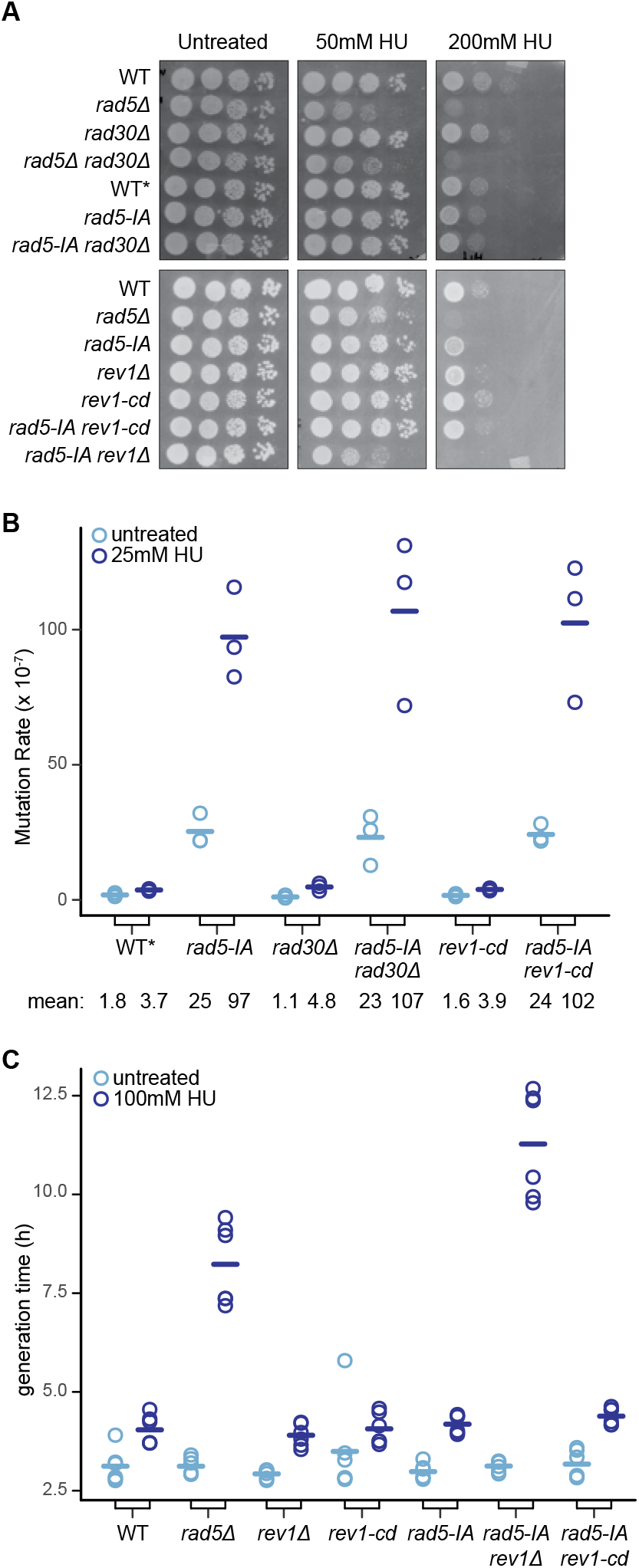
Rad5-mediated error-prone repair does not require Rad30 or the dCMP transferase activity of Rev1. (A) HU sensitivity assay of the indicated strains. Cultures were serially diluted and spotted on solid media containing the indicated concentrations of HU. Plates were incubated for three days prior to imaging. WT* (DGY19) carries *RAD5* tagged with 6His10FLAG, as do the *rad5-IA* strains. (B) Spontaneous and HU-induced mutation rates of the indicated strains are plotted. Each replicate (n = 3) is plotted, and solid bars represent the means, which are also indicated below the plot. WT* (DGY19) carries *RAD5* tagged with 6His10FLAG, as do the *rad5-IA* strains. Mutation rate data for WT* and *rad5-IA* are re-plotted from Figure 1C for ease of comparison. (C) Generation times of the indicated strains during logarithmic growth in the presence or absence of HU are plotted. Each replicate (n = 6) is plotted, and solid bars represent the means.

### Two modes of error-prone repair contribute to mutagenesis in the absence of error-free repair

In PRR, the canonical mutagenic repair pathway is mediated by the interaction between PCNA-Ub and the ubiquitin-binding motifs (UBM) of Rev1 (Guo et al. 2006; Wood et al. 2007). Mutating the conserved amino acids L821, P822, I825 in the UBM2 domain of Rev1 to alanines disrupts the PCNA-Rev1 interaction *in vitro* and abolishes PCNA-Ub-dependent damage-induced mutagenesis (Wood et al. 2007). To test whether the interaction between Rev1 and PCNA-Ub is required for mutagenic repair in *rad5-IA*, we introduced three mutations (L821A, P822A, I825A) into *REV1* to yield *rev1-LPI* (Figure 1A). The *rad5-IA rev1-LPI* strain has increased HU sensitivity relative to *rad5-IA* (Figure 3A), suggesting that the canonical recruitment of Rev1 by PCNA-Ub is required for mutagenic repair in *rad5-IA*. Intriguingly, *rad5-IA rev1-LPI* is not as sensitive to HU as *rad5-IA rev1Δ* (Figure 3A), indicating that there are additional modes of Rev1 recruitment in *rad5-IA* that are distinct from PCNA-Ub. Indeed, using a F13A, N14A mutant of Rad5 (*rad5-FNAA*) (Figure 1A) which disrupts the interaction between Rad5 and Rev1 (Xu et al. 2016), we observed an additive increase in HU sensitivity in *rad5-IA-FNAA rev1-LPI*, which is defective in both Rad5-mediated and PCNA-Ub-mediated recruitment of Rev1 (Figure 3A). These results suggest to us that Rad5-mediated mutagenic repair and PCNA-Ub-mediated mutagenic repair are separable, and that both pathways contribute to replication stress response in *rad5-IA*.

**Figure 3.**
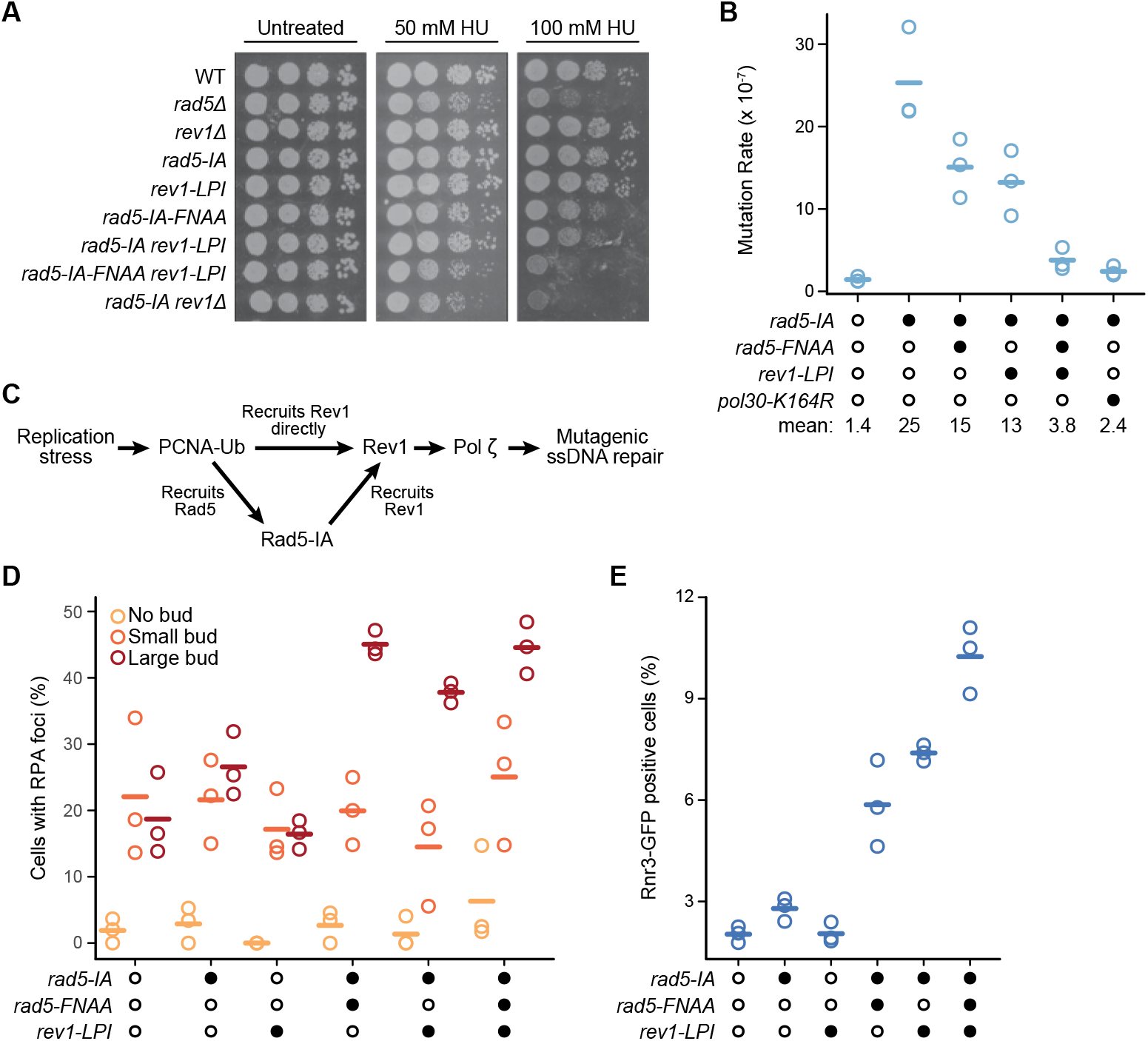
Two modes of error-prone repair contribute additively and independently to mutagenesis in *rad5-IA*. (A) HU sensitivity assay of the indicated strains. Cultures were serially diluted and spotted on solid media containing the indicated concentrations of HU. Plates were incubated for three days prior to imaging. (B) Spontaneous mutation rates of the indicated strains are plotted. Filled circles indicate the genotype of each strain, with wild type in the left column. Each replicate (n = 3) is plotted, and solid bars represent the means, which are also indicated below the plot. (C) Proposed model of the two error-prone repair pathways in *rad5-IA*. (D) Quantification of RPA focus formation in the indicated strains. Filled circles indicate the genotype of each strain, with wild type in the left column. At least 100 cells were counted for each strain, grouped based on the budding status, and the percentage of cells with RPA foci is plotted. Each replicate (n = 3) is plotted, and solid bars represent the means. (E) Percentage of cells expressing Rnr3-GFP is plotted. Filled circles indicate the genotype of each strain, with wild type in the left column. A total of 10,000 events was quantified by flow cytometry for each replicate, each replicate (n = 3) is plotted, and solid bars represent the means.

To further evaluate the contributions of Rad5-mediated and PCNA-Ub-mediated mutagenic repair in *rad5-IA*, we measured the spontaneous mutation rates of the individual strains. As expected, both *rad5-IA-FNAA* and *rad5-IA rev1-LPI* have decreased mutation rates relative to *rad5-IA*, indicating that mutagenic repair is compromised in these strains (Figure 3B, p = 0.06 and p = 0.04). Importantly, we observed an additive decrease of mutation rate in *rad5-IA-FNAA rev1-LPI* that reached a level similar to the spontaneous mutation rate of wild-type (Figure 3B, p = 0.007 and 0.02 when comparing to the double mutants). Finally, we measured the mutation rate of *rad5-IA pol30-K164R*, which cannot be post-translationally modified at K164 of PCNA (Hoege et al. 2002). The *pol30-K164R* mutation abolished mutagenesis in *rad5-IA*, suggesting that both modes of mutagenic repair act downstream of PCNA-Ub (Figure 3B). Based on these data, we propose that when the error-free template switching function of Rad5 is compromised, mutagenic repair of ssDNA gaps by Pol ζ occurs through two distinct pathways downstream of PCNA-Ub. In the canonical pathway, PCNA-Ub directly recruits Rev1. In the Rad5-mediated pathway PCNA-Ub first recruits Rad5 which then recruits Rev1 (Figure 3C). The *rev1-LPI* mutant disrupts the PCNA-Ub-mediated mutagenic repair, whereas the *rad5-FNAA* mutant disrupts the Rad5-mediated mutagenic repair.

### Disruption of either mode of mutagenic repair leads to ssDNA accumulation and checkpoint activation

We next asked whether both modes of mutagenic repair are required to prevent ssDNA accumulation. Both *rad5-IA-FNAA* and *rad5-IA rev1-LPI* showed higher percentage of large-budded cells with RPA foci compared to *rad5-IA*, indicating that losing either Rad5- or PCNA-Ub-mediated mutagenic repair results in ssDNA accumulation (Figure 3D, p = 0.004 and 0.02). However, while *rad5-IA-FNAA rev1-LPI* also has increased percentage of large-budded cells with RPA foci, no additive relationship was observed between the mutagenic repair pathway mutants (Figure 3D). We postulated that this was due to the dynamic range of spontaneous Rfa1 foci in PRR mutants, as the percentage of large-budded cells with RPA foci plateaus at around 50% even for *rad5-IA rev3Δ* (Figure 1D).

To test whether the two mutagenic repair pathways additively contribute to endogenous replication stress response in other aspects, we evaluated the expression of Rnr3. In budding yeast, Rnr3 expression is induced when the replication stress check-point is activated (Huang et al. 1998; Mulder et al. 2005). Rnr3 expression has been used as a reporter for DNA damage (Zhou and Elledge 1992; Wei et al. 2013; Hendry et al. 2015; Suzuki et al. 2017), and deletion of *RAD5* induces Rnr3 expression (Hendry et al. 2015; Gallo et al. 2019). We found that both *rad5-IA-FNAA* and *rad5-IA rev1-LPI* have more cells expressing Rnr3 than does *rad5-IA* (Figure 3E, p = 0.02 and 0.00005), and the number of Rnr3-positive cells is further increased in *rad5-IA-FNAA rev1-LPI* (Figure 3E, p = 0.009 and 0.009 when comparing to the double mutants), indicating an additive effect on checkpoint activation when both modes of mutagenic repair are defective.

### Rad5- and PCNA-Ub-mediated mutagenic repair act independent of replication timing

Genomic regions with late replication timing are associated with higher mutation rate (Agier and Fischer 2012; Supek and Lehner 2015). This phenomenon is linked to error-prone repair by Rev1-Pol ζ (Lang and Murray 2011; Seplyarskiy et al. 2015), since deletion of *REV1* abolishes mutagenesis in late-replicating regions and the Pol ζ mutation signature is associated with late replication timing in humans. Given that we dissected two mutagenic repair pathways in *rad5-IA*, we asked whether the different pathways account for mutation rate variation at regions with different replication timing. To test this, we used strains with *URA3* replacing non-essential open reading frames located at early- or late-replicating regions on chromosome VI (Lang and Murray 2011). In agreement with the previous analysis, strains with *URA3* at late-replicating regions showed higher *URA3* mutation rate than those at early-replicating regions, and the difference in mutation rates is dependent on *REV1* (Lang and Murray 2011) (Figure 4A). Introducing *rad5-IA* into the *URA3* strains led to increased mutation rates in all strains tested, regardless of the position of *URA3* (Figure 4B).

We reasoned that if Rad5-mediated or PCNA-Ub-mediated mutagenic repair act preferentially at either early- or late-replicating regions, disruption of one mutagenic repair pathway should reduce the *rad5-IA* mutagenesis in either early or late-replicating *URA3* but not in both. When *rad5-FNAA* or *rev1-LPI* were introduced with *rad5-IA*, there was a clear trend of decreased *URA3* mutagenesis at all regions, although strong statistical support for decreased mutagenesis was lacking for two comparisons (Figure 4B; p = 0.08 for rev1-LPI 26.5 min, p = 0.07 for rad5-FNAA 43.8 min, all others p ≤ 0.05). Since early- and late-replicating regions were both affected similarly by disruption of either mutagenic repair pathway, we conclude that both modes of mutagenic repair contribute to repair at early- and late-replicating regions.

### Rad5- and PCNA-Ub-mediated mutagenic repair contribute to mutagenesis in a natural RAD5 variant

Mutation rate varies among *S. cerevisiae* strains, and the *RAD5* locus has been associated with mutation rate variation (Gou et al. 2019). The *RAD5* gene in the vineyard strain RM11-1a encodes amino acid substitutions (E783D and I791S) that are absent in the laboratory strain BY, and that account in part for the higher mutation rate of RM11-1a (Gou et al. 2019). We hypothesized that the increased mutation rate in RM11-1a could be due to the activities of the Rad5- and PCNA-Ub-mediated mutagenic repair pathways that we defined. To test this hypothesis, we introduced the E783D and I791S mutations (Figure 5A, hereafter refer to as *rad5-RM*) into our laboratory BY4741 strain, which led to an increase in HU sensitivity and a large increase in mutation rate (Figure 5B and 5C, p = 0.001). Importantly, disrupting either mode of mutagenic repair with *rad5-FNAA* or *rev1-LPI* led to an increase in HU sensitivity that appeared additive in the *rad5-RM-FNAA rev1-LPI* triple mutant (Figure 5B). The *rad5-RM-FNAA* and *rad5- RM rev1-LPI* mutants also have decreased mutation rates relative to *rad5-RM* (Figure 5C, p = 0.02 and 0.003), and we saw a further decrease in mutation rate for *rad5-RM-FNAA rev1-LPI* (Figure 5C, p = 0.05 and 0.02 when compared to the double mutants). These data are reminiscent of *rad5-IA* (Figure 3A and 3B), and so suggest that the two modes of mutagenic repair, mediated by Rad5 and PCNA-Ub, contribute to mutagenesis in RM11-1a and to mutation rate variation in wild strains.

**Figure 5.**
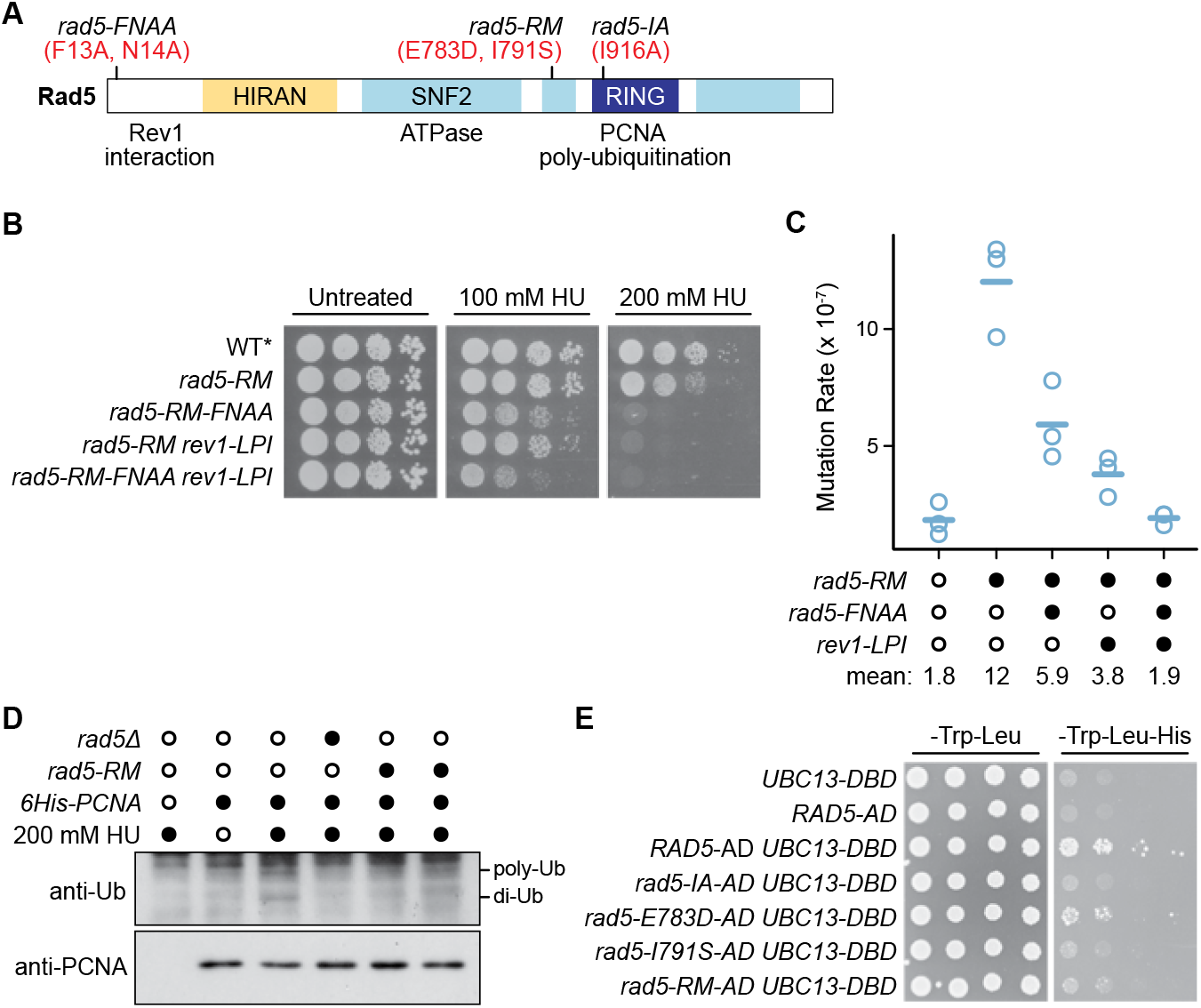
A RAD5 natural variant is defective in PCNA poly-ubiquitination. (A) Schematic of Rad5 showing the positions of the amino acid substitutions encoded by *rad5-IA* and *rad5-RM*. (B) HU sensitivity assay of the indicated strains. Cultures were serially diluted and spotted on solid media containing the indicated concentrations of HU. Plates were incubated for three days prior to imaging. WT* (DGY19) carries *RAD5* tagged with 6His10FLAG, as do the *rad5-RM* strains. (C) Spontaneous mutation rates of *rad5-RM* strains are plotted. Filled circles indicate the genotype of each strain, with wild type (WT*) in the left column. Each replicate (n = 3) is plotted, and solid bars represent the means, which are also indicated below the plot. WT* (DGY19) carries *RAD5* tagged with 6His10FLAG, as do the *rad5-RM* strains. Mutation rate data for WT* are re-plotted from Figure 1C for ease of comparison. (D) PCNA ubiquitination assessed by immunoblotting. His-tagged PCNA was affinity-purified under denaturing conditions from the indicated strains, following growth in 200 mM HU. The wild-type strain lacking tagged PCNA is included as a control, as is wild-type with tagged PCNA without HU treatment. Two biological replicates are shown for *rad5-RM*. (E) Yeast 2-hybrid analysis of protein interactions between activation domain (AD)-tagged Rad5 variants and DNA binding domain (DBD)-tagged Ubc13. Cultures of strains carrying the indicated AD and DBD plasmids were serially diluted and spotted on SD-Trp-Leu or SD-Trp-Leu-His solid media. Growth on SD-Trp-Leu-His indicates a protein-protein interaction. Plates were incubated for three days prior to imaging.

The amino acid substitutions encoded by *rad5-IA* lie in the RING domain of Rad5 and disrupt PCNA poly-ubiquitination by perturbing the interaction between Rad5 and Ubc13 (Ulrich 2003; Toth et al. 2022). Given the heightened activity of mutagenic repair in *rad5-RM*, we tested if poly-ubiquitination of PCNA is compromised in *rad5-RM*. We detected PCNA ubiquitination by affinity purifying PCNA under denaturing conditions and probing for ubiquitin by immunoblotting (Davies and Ulrich 2012). We found that HU-dependent di-ubiquitination, which requires Rad5, was reduced in two independent isolates of *rad5-RM* (Figure 5D). We then took a yeast 2-hybrid (Fields and Song 1989) approach to determine whether the physical interaction between Rad5 and Ubc13 was impaired (Figure 5E). Rad5-RM showed a profound defect in binding to Ubc13, similar to that observed with Rad5-IA. We separated the two amino acid substitutions in Rad5-RM and found that the defective interaction with Ubc13 was mainly due to I791S, while E783D displayed only a modest interaction defect. Based on these data we infer that *rad5-RM* is defective in PCNA poly-ubiquitination due to decreased interaction between Rad5-RM and Ubc13, resulting in reduced error-free repair and increased mutagenic repair.

## Discussion

We used genetic approaches to define two mutagenic repair pathways that function when the template switching activity of Rad5 is compromised. In our model (Figure 6), ssDNA arises due to replication stress and PCNA is mono-ubiquitinated, which can either recruit Rad5 for Rad5-mediated mutagenic repair, or directly recruit Rev1 for the canonical PCNA-Ub-mediated mutagenic repair. Both pathways require the TLS DNA polymerase ζ, and defects in either mutagenic repair pathway lead to accumulation of ssDNA gaps and to checkpoint activation, even in otherwise unperturbed conditions. Finally, we showed that both mutagenic repair pathways contribute to mutation rate variation in a *S. cerevisiae* wild strain.

**Figure 6.**
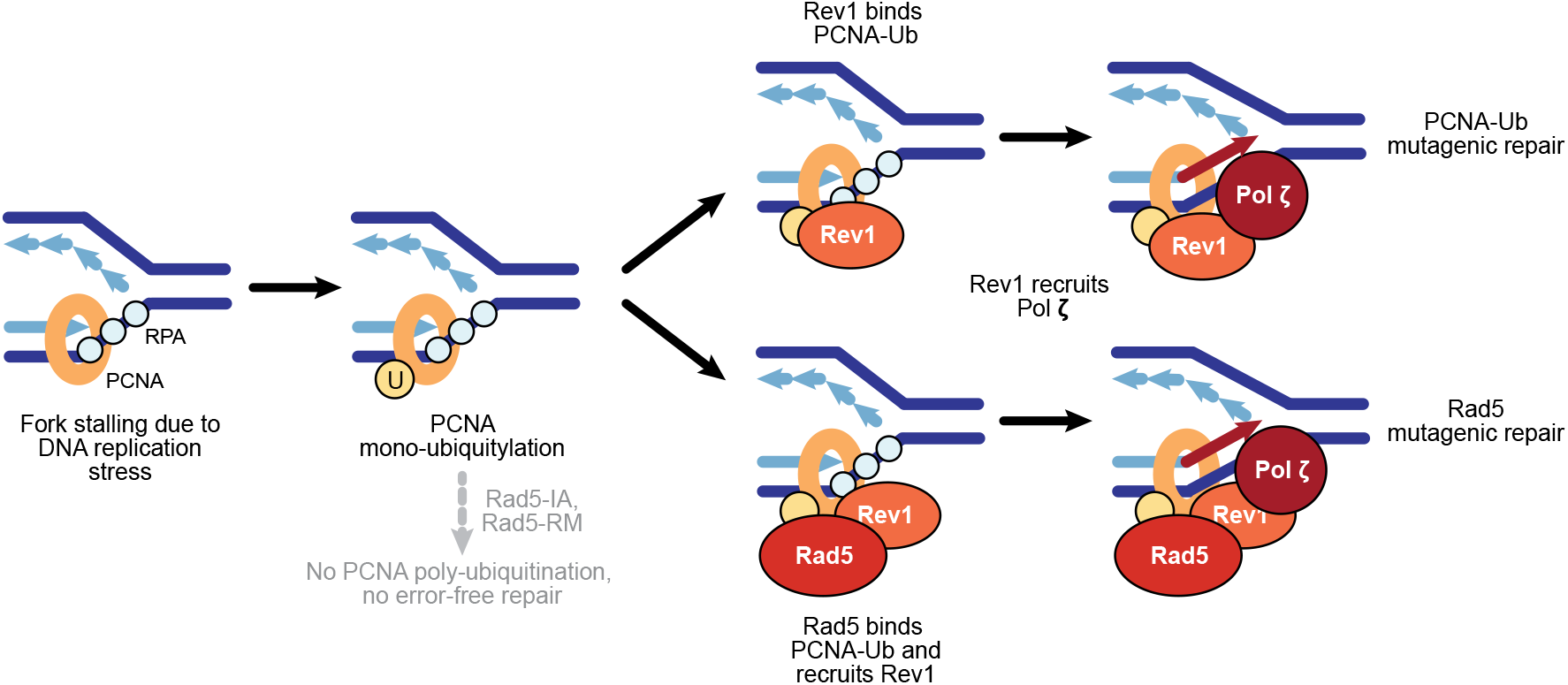
Model of two mutagenic repair pathways in template switching defective Rad5 mutants. During DNA replication stress, PCNA-Ub can directly recruit Rev1 via the ubiquitin binding motifs of Rev1, or recruit Rad5. Rad5 brings Rev1 to the ssDNA repair site via interaction of the Rad5 amino-terminal domain with Rev1. Rev1 acts as a scaffold for the TLS polymerase Pol ζ, which repairs the ssDNA independent of base insertion by Rev1.

### ssDNA resulting from DNA replication stress is repaired by Pol ζ

We tested whether each of the three yeast TLS polymerases participates in mutagenic repair when error-free repair is compromised by the *rad5-IA* mutation, revealing that Pol ζ activity and a non-catalytic function of Rev1 are required. The lack of requirement for Rad30/Pol η agrees with previous findings that Rad30 is dispensable in PRR of UV-induced lesions (Daigaku et al. 2010) and for spontaneous mutagenesis induced by DNA replication stress in *mcm10-1* mutants (Becker et al. 2014). Although the dCMP transferase activity of Rev1 is involved in bypassing short hairpin structures that form due to replication stress in Pol δ-deficient strains (Northam et al. 2014), we did not observe a contribution of Rev1 catalytic activity to the replication stress response in *rad5-IA*. Since *rev1Δ* was defective in mutagenic repair, we infer that it is a structural role of Rev1 that is important.

The protein-protein interactions of Rev1 are consistent with our experimental data and provide a reasonable explanation for the function of Rev1 in each mutagenic repair pathway. In the case of PCNA-Ub-mediated mutagenic repair, the ubiquitin-interacting motifs of Rev1 are required to recruit Pol ζ to mono-ubiquitinated PCNA (Guo et al. 2006; Wood et al. 2007). In the case of Rad5-mediated pathway, the Rev1 binding site on Rad5 is important for recruiting Pol ζ to Rad5 engaged at replication stress sites (Xu et al. 2016; Gallo et al. 2019).

### Mutagenic repair downstream of DNA replication stress requires the catalytic activity of Pol ζ

Pol ζ is the only TLS polymerase that belongs to the B-family of DNA polymerases, which includes the replicative polymerases ε and δ. High accuracy and processivity characterize the B-family polymerases (McCulloch and Kunkel 2008; Yang 2014) in comparison to Y-family polymerases. Pol ζ typically functions as an extender polymerase in TLS (Johnson et al. 2000; Haracska et al. 2003; Lee et al. 2014; Bezalel-Buch et al. 2020), replicating up to 1 kb downstream of UV lesions for example (Kochenova et al. 2015). Pol ζ is less accurate than the other B-family polymerases as it lacks proof-reading activity (Makarova and Burgers 2015) and has lower nucleotide selectivity (Arana and Kunkel 2010). Yeast Pol ζ is recruited by polymerase stalling at short hairpins and can mediate both insertion and extension (Northam et al. 2014), but the details of how Pol ζ is engaged at undamaged daughter strand gaps are unclear. Our data indicate that recruitment of Pol ζ by Rev1 through binding to Rad5 or to PCNA-Ub is a key element. Given that Pol ζ readily extends terminal mismatched nucleotides (Johnson et al. 2000; Haracska et al. 2003), and that DNA replication stress in the form of dNTP pool perturbations increases mismatched nucleotide incorporation (Kumar et al. 2011; Watt et al. 2016), one possibility is that Pol ζ is engaged when replicative polymerases fail to extend mismatches.

We describe the presence of two mutagenic repair pathways contributing to ssDNA gap repair when the template switching activity of Rad5 is compromised. Both of the pathways act down-stream of PCNA-Ub, in agreement with our previous observation that PCNA-Ub is required to recruit Rad5 to HU-stalled replication forks (Gallo et al. 2019). Importantly, both modes of mutagenic repair are required for a robust replication stress response, as the disruption of either pathway leads to ssDNA accumulation and checkpoint activation. We find that both mutagenic repair pathways depend on the polymerase activity of Pol ζ, indicating that although two distinct recruitment modes exist, the repair events themselves are likely to be similar. In addition, both pathways require Rev1, but not Rev1 catalysis, indicating that mutagenic repair events in the absence of *RAD5* error-free repair are not initiated by Rev1 inserter activity. In a similar vein, Rev1 catalytic activity is not important for defective-replisome-induced mutagenesis in a *mms2Δ* background, where error-free repair is absent (Northam et al. 2014). Of particular interest, the *REV1* role in mutagenic repair in HU-induced replication stress mirrors the *REV1* role in mutagenic repair in unperturbed cells (structural and not catalytic), suggesting that endogenous replication stress is also processed by Pol ζ without a preceding base insertion by Rev1.

### The mutagenic repair pathways both act at early and late replicating regions

Mutation rate in *S. cerevisiae* varies according to chromosomal location, and associates with replication timing, with late replicating regions having higher mutation rates (Lang and Murray 2011; Sharp et al. 2018). In metazoans the association of mutagenesis with late replication is also evident, particularly in cancer genomes (Woo and Li 2012; Liu et al. 2013), and has been attributed to more effective mismatch repair at early replicating regions (Supek and Lehner 2015). Surprisingly, the higher mutation rates at late replicating regions in yeast depend on Rev1 (Lang and Murray 2011), even though interactions between Rev1 and chromatin favor regions proximal to early-firing replication origins over late-firing origins during DNA replication stress (Gallo et al. 2019). Rad5 also appears to be subject to temporal regulation, although the exact nature of this remains unclear. During DNA replication stress in early S phase Rad5 is strongly associated with replication fork proximal regions (Gallo et al. 2019). However, in the presence of template strand lesions, post-replication repair promoted by Rad5 can certainly occur on daughter strand gaps that are distal to replication forks (Daigaku et al. 2010; García-Rodríguez et al. 2018; Wong et al. 2020). We tested whether Rad5- and PCNA-Ub-dependent mutagenic repair might act differentially at early-vs late-replicating region and found that both pathways act irrespective of DNA replica-tion timing. We conclude that the Rev1-dependent mutagenesis bias in replication timing is not due to different modes of Rev1 recruitment, but could be due to other factors such as increased Rev1 protein abundance in G2 phase (Waters and Walker 2006). Alternatively, the two mutagenic repair pathways might be differentiated by other aspects, such as their preferred cell cycle stage or whether they act at the fork or distal to the fork. A recent study in cisplatin-treated human cells showed that REV1-POL ζ repairs PRIMPOL-dependent ssDNA gaps in both S-phase and G2-phase, whereas error-free template switching factors UBC13 and RAD51 can only repair ssDNA gaps in S phase (Tirman et al. 2021). Given that the protein levels of Rad5 and Rev1 peak at different stages of cell cycle (Waters and Walker 2006; Ortiz-Bazán et al. 2014), it is possible that the two mutagenic pathways repair ssDNA in a temporally different manner. Of course, it is also possible that the two pathways act in concert in all contexts to ensure rapid gap-filling and timely completion of DNA replication in a truly redundant fashion.

### The DNA helicase domain of Rad5 is important for RING domain function

Intriguingly, the *rad5-RM* polymorphisms are in the helicase domain of Rad5, rather than the RING domain where ubiquitin ligation is promoted. Although it is not known whether *rad5-RM* is defective in helicase activity, the lack of increased spontaneous mutation rate in the helicase-dead separation-of-function *rad5-Q1106D* mutant argues against a role for helicase catalytic activity in regulating the RING domain (Choi et al. 2015; Gallo et al. 2019). By contrast, *rad5-D681A,E682A*, which is a helicase domain mutant that is defective in both helicase activity and poly-ubiquitination activity, was unable to bind Ubc13, indicating that both the helicase domain and the RING domain are important for the Rad5-Ubc13 interaction (Choi et al. 2015). Finally, the RING domain of Rad5 is necessary but not sufficient to support Rad5 binding to Ubc13 (Ulrich and Jentsch 2000). Since Rad5-RM, like Rad5-D681A,E682A, is located in the helicase domain and perturbs the Rad5-Ubc13 interaction, we suggest that the helicase domain of Rad5 plays a structural role in maintaining the RING domain so that it can bind to Ubc13.

### Mutagenic repair pathways can cause natural variation in mutation rates

We exploited an engineered mutation in Rad5 (*rad5-IA*) to eliminate error-free repair, allowing us to establish the presence of two error-prone repair pathways. Our experimental context was, of course, artificial, and the comparatively low mutation rate of the parental strain with intact error-free repair indicates that in the BY strain background error-free repair predominates. Mutation rate in different species, and even in different strains of the same species, can vary by at least an order of magnitude (Fijarczyk et al. 2021; Jiang et al. 2021; Melde et al. 2022), and in most cases the mechanism for the variation is not understood. *RAD5* was identified as the main causal locus for the mutation rate variation between the RM11-1a strain and the BY strain (Gou et al. 2019), and we found that the *rad5* allele in RM11-1a confers increased mutagenesis catalyzed by both Rad5- and PCNA-Ub-mediated error-prone repair. In mechanistic terms, the mutation rate variation between RM11-1a and BY is due to reduced physical interaction between Rad5-RM and the E2 protein Ubc13 compromising PCNA polyubiquitination, thereby inactivating error-free repair of ssDNA gaps resulting from DNA replication stress. Thus, Rad5-mediated mutagenic repair exists in the wild, contributes to mutation rate variation, and generates genetic diversity.

## Supporting information

Reagent Table

## Acknowledgments

We thank Dr. Elçin Ünal for the CRISPR-Cas9 plasmid and protocol, Dr. Helle Ulrich for the YIplac128-His6-POL30 plasmid, Dr. Marco Muzi-Falconi for the PCNA antibody, Dr. Andrew Murray for the *URA3* reporter strains, Dr. Fritz Roth for the pDEST-AD and pDEST-DB Y2H vectors, and Dr. Charlie Boone for the Y8800 and Y8930 strains. We thank members of the Brown lab for careful reading of the manuscript, and helpful discussions. We are grateful to work on the lands of the Mississaugas of the Credit, the Anishnaabeg, the Haudenosaunee and the Wendat peoples, land that is now home to many diverse First Nations, Inuit, and Métis peoples.

## Author contributions

YKJ: Designed and carried out the experiments and data analysis, wrote the paper, edited the paper.

EM: Constructed strains, edited the paper.

GWB: Designed the experiments, wrote the paper, edited the paper.

