## Supplemental Figure S1 for "Two independent DNA repair pathways cause mutagenesis in template switching deficient *Saccharomyces cerevisiae*"

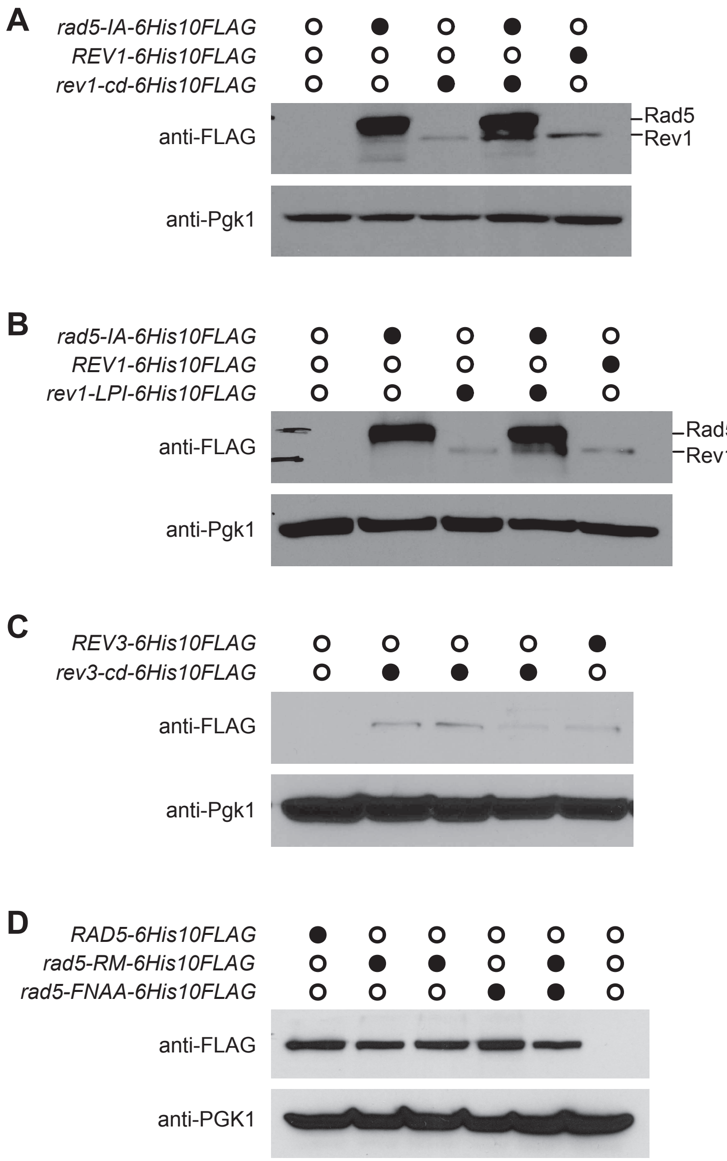


**Figure S1** Immunoblot analysis of strains expressing tagged Rad5, Rev1, and Rev3 proteins. (A – D) Extracts of strains expressing the indicated genes tagged with 6His10FLAG sequences were fractionated on SDS-PAGE and immunoblotted with anti-FLAG or anti-Pgk1 antibodies.
